# Capillary Electrophoresis of PCR fragments with 5’-labelled primers for testing the SARS-Cov-2

**DOI:** 10.1101/2020.05.16.099242

**Authors:** Juan Gómez, Santiago Melón, José A. Boga, Marta E. Alvarez-Argüelles, Susana Rojo-Alba, Alvaro Leal-Negredo, Cristian Castello-Abietar, Victoria Alvarez, Elías Cuesta-Llavona, Eliecer Coto

## Abstract

**Background:** Due to the huge demand for SARS-Cov-2 determination, alternatives to the standard qtPCR tests are potentially useful for increasing the number of samples screened. Our aim was to develop a direct fluorescent PCR capillary-electrophoresis detection of the viral genome. We validated this approach on several SARS-Cov-2 positive and negative samples.

**Study design:** We isolated the naso-pharingeal RNA from 20 positive and 10 negative samples. The cDNA was synthesised and two fragments of the SARS-Cov-2 were amplified. One of the primers for each pair was 5’-end fluorochrome labelled. The amplifications were subjected to capillary electrophoresis in ABI3130 sequencers to visualize the fluorescent peaks.

**Results:** The two SARS-Cov-2 fragments were successfully amplified in the positive samples, while the negative samples did not render fluorescent peaks.

**Conclusion:** We describe and alternative method to identify the SARS-Cov-2 genome that could be scaled to the analysis of approximately 100 samples in less than 5 hours. By combining a standard PCR with capillary electrophoresis our approach would overcome the limits imposed to many labs by the qtPCR (lack of reactive and real-time PCR equipment) and increase the testing capacity.

## Introduction

PCR tests for the accurate and fast SARS-CoV-2 testing are crucial for the identification of carriers and control the spreading of COVID19. Most of these tests rely on the isolation and retro-transcription of the viral RNA from nasal, pharyngeal or pulmonary specimens followed by real-time quantitative PCR (qtPCR) with primers/probes designated from the SARS-Cov-2 sequence [1–4]. Most of the labs use commercial kits that are usually not supplied on demand due to the huge global demand. This is a bottle-neck in massive screening of this new coronavirus, and reinforces the necessity of developing new technical approaches to overcome these barriers. We describe a technical approach to SARS-Cov-2 testing by amplifying fragments of the viral genome with 5’-fluorescent primers followed by capillary electrophoresis in an ABI3130xl equipment. This method permits the analysis of 96 samples in approximately five hours.

## Methods

### Samples preparation

Our method was validated in nasal-swap samples from 30 individuals, 20 positive and 10 negative for SARS-Cov-2 testing with a standard qtPCR. The total RNA was isolated with MagNa Pure-96 System (Roche) in a final vol of 100 μL. A total of 10 μL of the RNA were retrotranscribed (RT) with the RetroScript™ kit (*Invitrogene*) in a final vol of 20 μL. To check for quality of the cDNA synthesis we amplified a fragment of the human beta-actin gene (*ACTB*) and two fragments of the SARS-Cov-2. Briefly, two μL of the RT reactions were subjected to 32 PCR cycles with primer-pairs specific for either *ACTB* or the viral genome in a final vol of 30 μL. Ten μL of the PCRs were electrophoresed on 3% agarose gels to visualize the presence of the corresponding fragments. The 30 samples rendered an *ACTB* fragment, and we thus validated the 30 samples for the presence of cDNA. The *ACTB* transcript was also confirmed in all the samples by a qtPCR (**supplementary figure 1**). The 20 positive samples amplified the two SARS-Cov-2 fragments, that were not visualised in the 10 negative samples (**Figure 1**). The viral fragments from the 20 positive samples were Sanger sequenced with *BigDye* chemistry in an ABI3130xl equipment (**supplementary figures 2, 3**). The SARS-Cov-2 primers were designated from the reference sequence in the NCBI Virus (https://www.ncbi.nlm.nih.gov/labs/virus/vssi/#/) [5].

**Figure 1.**
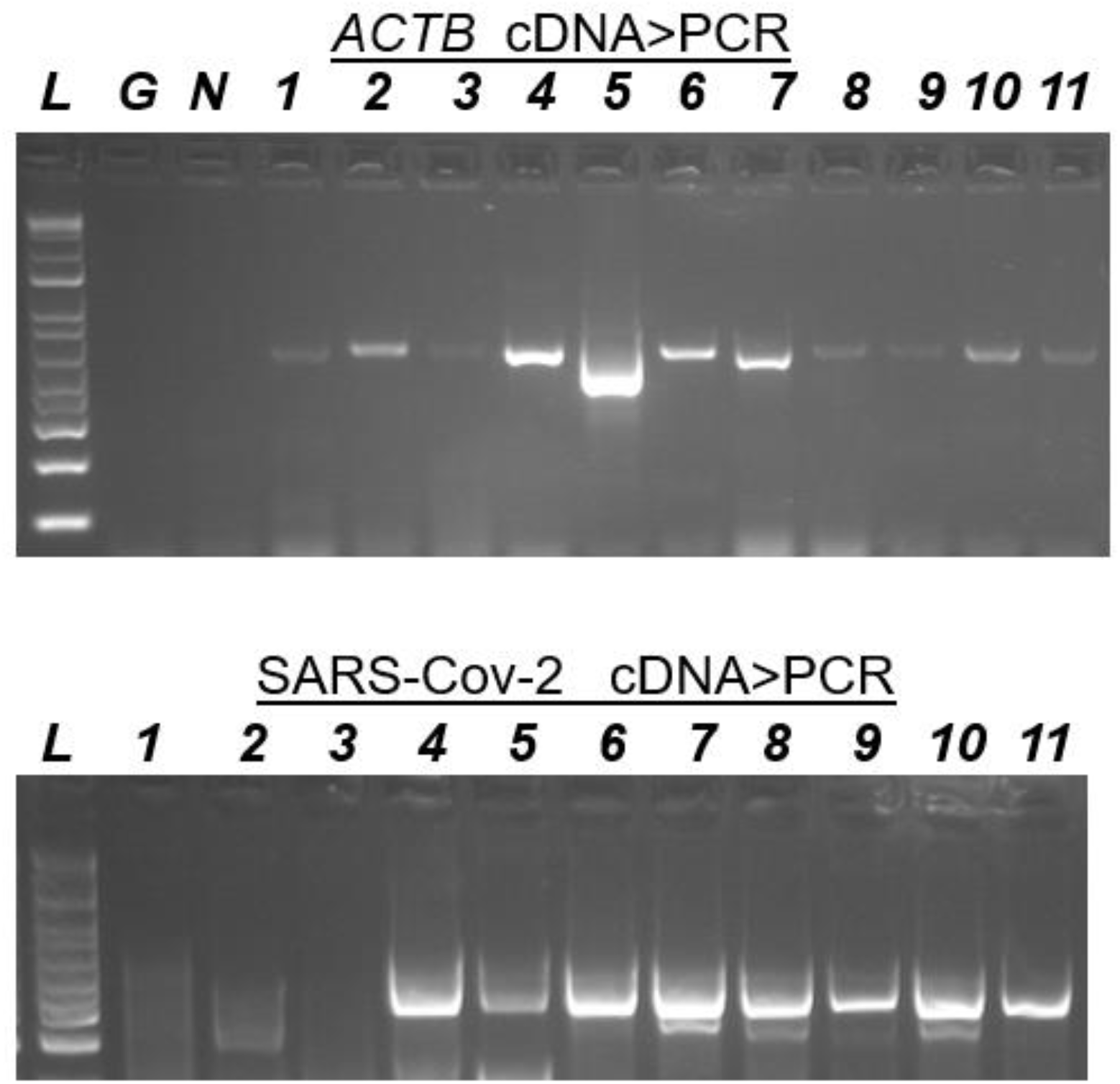
**Upper:** PCR fragments (agarose gels) for the human *ACTB* gene in cDNAs from nasal-samples (lanes 1-11). G and N, lack of amplification in the tubes containing genomic DNA and water (negative controls). **Lower:** PCR of a SARS-Cov-2 genome in cDNAs from viral positives (4-11) and negatives (1-3). These fragments were sanger sequenced (supplementary figures). L= 1 kb ladder, size marker.

### Fluorescent PCR and capillary electrophoresis

We designated two primer pairs to amplify fragments of the two viral genome Sanger sequenced fragments. To avoid the chance of false negatives the primers were specific for non-polymorphic sites in the 20 positive samples. The forward primers were 5’-end labelled with the 5-FAM or HEX fluorochromes. Two μL of the cDNAs were subjected to PCR with the two primer pairs in the same tube (30 μL final volume; 32 cycles of 95°C-30 s, 62°C-30s, 72°C-30s). The primers for fragment 1 were 5-FAM-5’CTTCACACTCAAAGGCGGTGCACC (forward) and 5’GCGAACTCATTTACTTCTGTACCGAGTTC (fragment size=180 bp). The primers for fragment 2 were HEX-5’GCTG TCAAATTACAGAATAATGAGCTTAGTCC (forward) and 5’CGGATAACA GTGCAAGTACAAACCTACC (fragment size=162 bp). Ten μL of the PCRs were mixed with 20 μL of deionised formamide and 0.5 μL of the fragment size-marker (250-ROX) in 96-well plates. The samples were subjected to capillary electrophoresis in an *ABI3130xl* and analysed with the Gene-Mapper software.

## Results and discussion

We processed 20 SARS-Cov-2 positive and 10 negative samples. The accuracy of the viral RNA isolation and RT synthesis was validated by amplifying a fragment of the human *ACTB* gene (**Figure 1**). In the 20 positive samples we amplified two SARS-Cov-2 fragments that were further sequenced to determine the presence of nucleotide changes (**Supplementary figures 2, 3**). We identified three patients who were heterozygous for 3,037 T/C and 8,782 T/C, while 17 were 3,037 T and 8,782 C. The two are nucleotide changes reported from viral isolates worldwide, and 8,782 C is in disequilibrium with *ORF8*: C251T (p.S84L in the spike protein) [6–9].

Based on the observed viral sequences we designated two primer pairs to amplify fragments of approximately 120 bp. The forward primers contained a 5’-end fluorochrome (5-FAM or HEX). The two were amplified in the same tube (multiplex PCR). The capillary electrophoresis of the fluorescent PCRs showed the two peaks in the SARS-Cov-2 samples, and its absence in the negatives (**Figure 2**).

**Figure 2.**
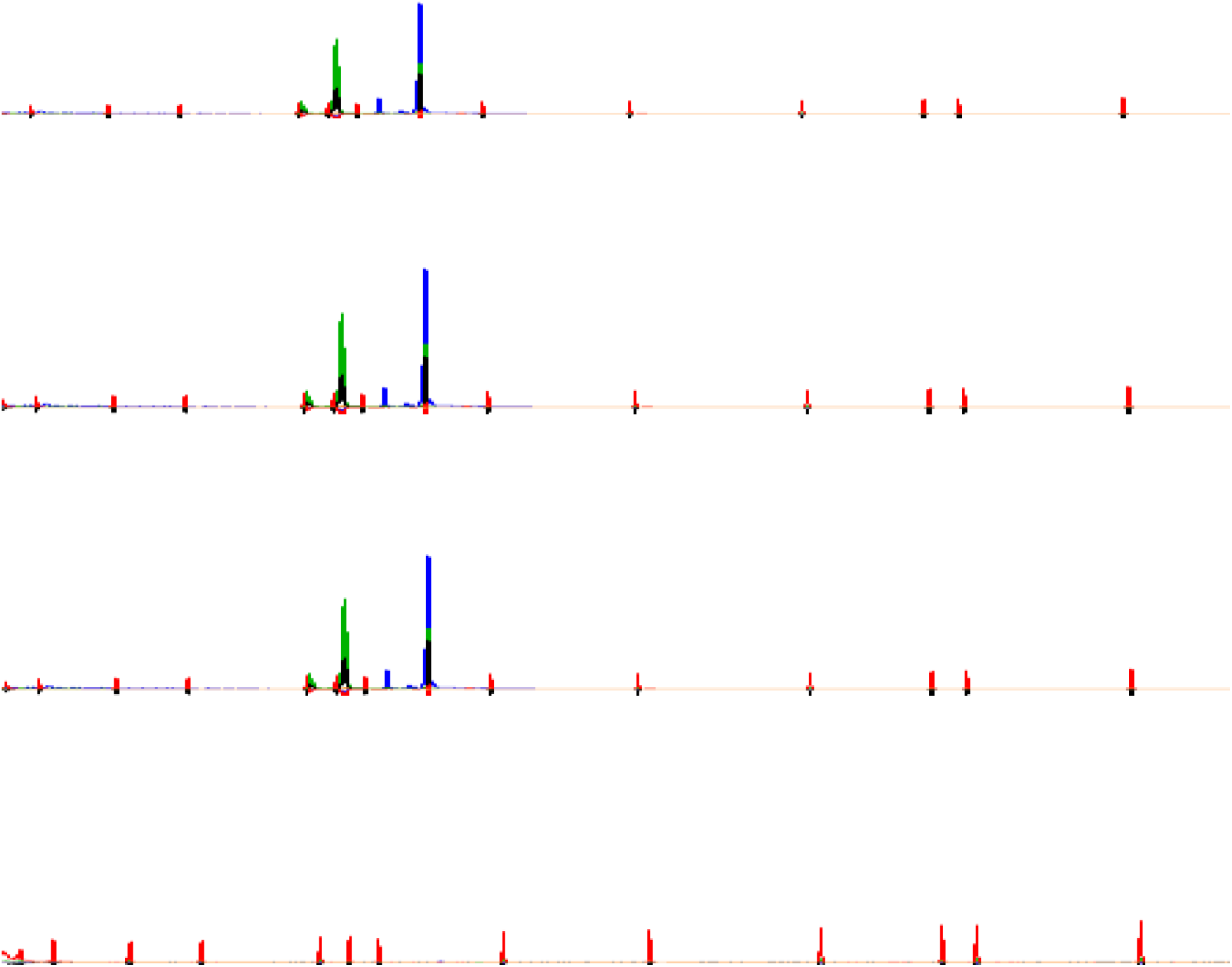
Capillary electrophoresis of the fluorescent PCR fragments from three SARS-Cov-2 positive cDNAs (three upper lines) and a negative sample (lower line). The two peaks corresponded to fragment 1 (blue, 180 bp) and fragment 2 (green, 162 bp). The red peaks are from the size-marker (250-ROX).

The total amplification time of our protocol takes about 90 min, and for 96-well plates the electrophoresis should be complete in approximately 3 hours. Thus, the analysis of 96 samples could be done in < 5 hours. The cDNA and PCR steps could be performed simultaneously with one-step RT and PCR kits with a reduction of the required time.

The protocol here described for the identification of SARS-Cov-2 is flexible in terms of the viral genome fragments prone to amplification, and could permit the simultaneous detection of multiple fragments (multiplex PCR). Moreover, compared to qtPCR the fluorescent PCR is not limited for the “compatibility” between the Taqman probe and the flanking PCR primers. An almost unlimited combination of primers to amplify the viral genome is thus possible. Also, because the fluorescent primers are purchased on demand for thousand of reactions and the other are common reactive with many suppliers there is not a foreseeable bottle-neck for the analysis of large amounts of samples. Our method should be particularly useful for virology labs that have capillary sequencers in their facilities, and could thus validate the new primer sets for capillary electrophoresis with the gold standard real-time PCR method.

In conclusion, we present an alternative method for the detection of the SARS-Cov-2 genome in biological samples. Our approach was based on capillary electrophoresis of fluorescent PCRs, and permits the screening of 96 samples in less than 5 hours. The method was validated with 20 virus positive and 10 virus negative samples, and would require a validation by each lab before its application.

## Contributorship

All the authors contributed to this work by recruiting the cohorts or performing the genetic and statistical analysis.

---

**Supplementary figure.**
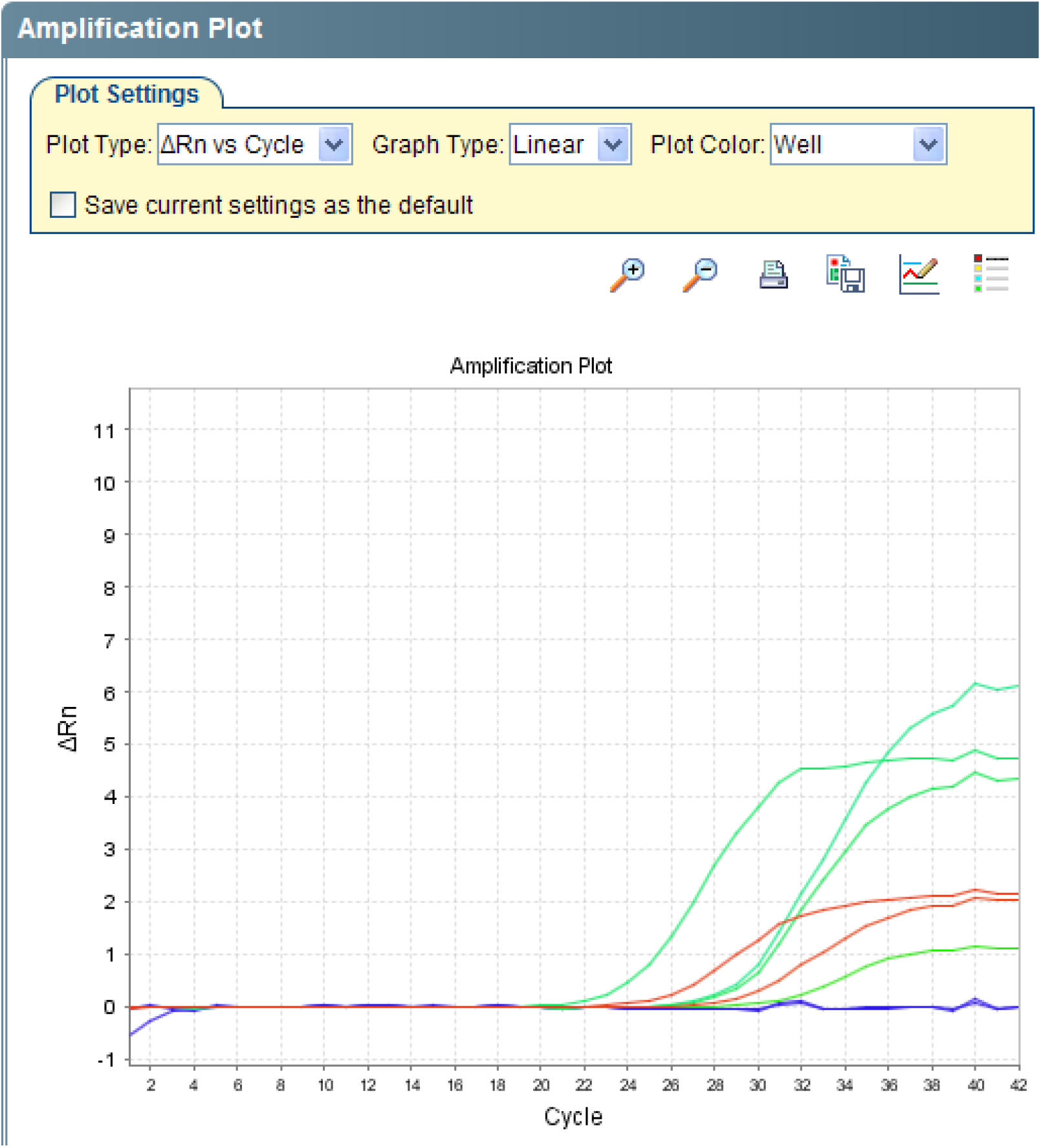
Real time quantitative PCR with Taqman assay for the human *ACTB* gene. A typical amplification profile was observed for the cDNAs from nasal-swaps in either viral positive (green) and negatives (red). Blue, negative control (genomic DNA in the PCR tube).

**Supplementary figure.**
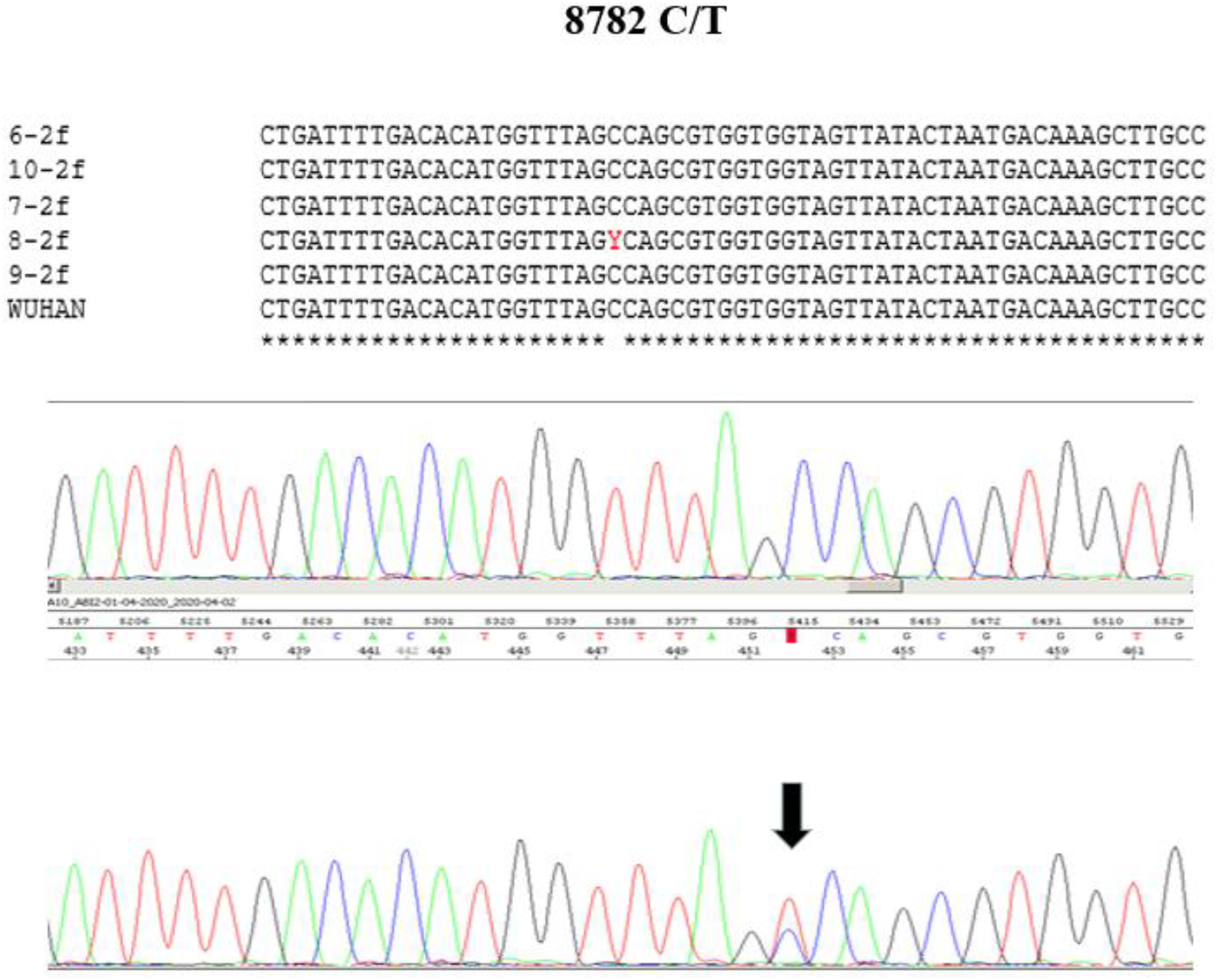
Sanger electropherogram of a SARS-Cov-2 PCR fragment generated with primers CV2F-CAT GAC ACC CCG TGA CCT TGG T and CV2R-CGT GCC AGG CAA ACC AGG CAC. Arrow shows the polymorphic site at nucleotide 8782 of the viral genome. The partial alignment of 5 cDNAs relative to the reference Wuhan genome is also shown.

**Supplementary figure.**
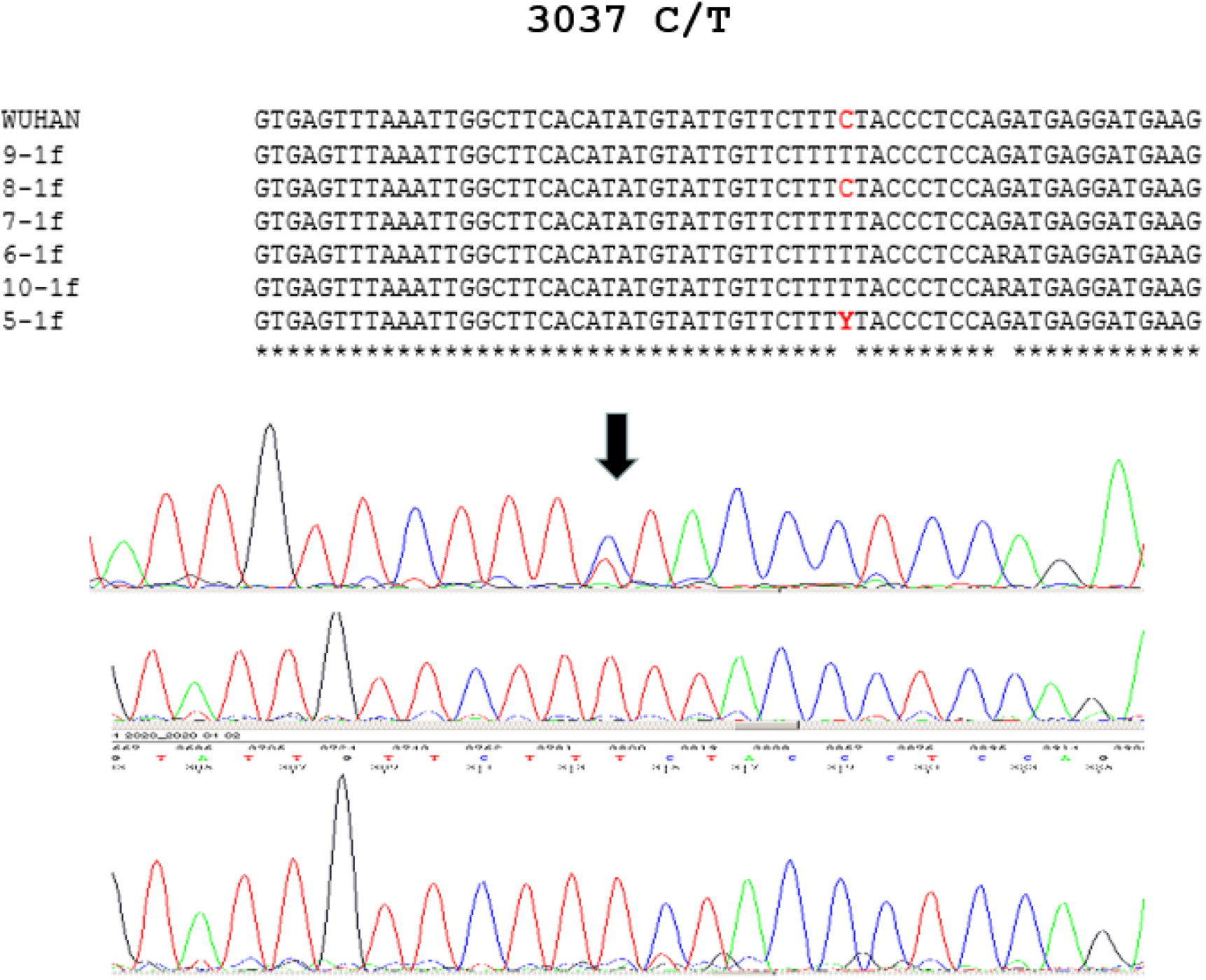
Sanger electropherogram of a SARS-Cov-2 PCR fragment generated with primers CV1F-CTT CAC ACT CAA AGG CGG TGC ACC and CV1R-CCA TCT CTA ATT GAG GTT GAA CCT CAA C. The arrow indicates the polymorphic site at nucleotide 3037 of the viral genome. The partial alignment of 6 cDNAs relative to the reference Wuhan genome is also shown.

